# A compendium of predicted growths and derived symbiotic relationships between 803 gut microbes in 13 different diets

**DOI:** 10.1101/2021.04.10.439264

**Authors:** Rohan Singh, Anirban Dutta, Tungadri Bose, Sharmila S. Mande

## Abstract

Gut health is intimately linked to dietary habits and the microbial community (microbiota) that flourishes within. The delicate dependency of the latter on nutritional availability is also strongly influenced by symbiotic relationships (such as, parasitic or mutualistic) between the resident microbes, often affecting their growth rate and ability to produce key metabolites. Since, cultivating the entire repertoire of gut microbes is an infeasible task, metabolic models (genome-based metabolic reconstructions) could be employed to predict their growth patterns and interactions. Here, we have used 803 gut microbial metabolic models from the Virtual Metabolic Human repository, and subsequently optimized and simulated them to grow on 13 dietary compositions. The presented pairwise interaction data (https://osf.io/ay8bq/) and the associated bacterial growth rates are expected to be useful for (a) deducing microbial association patterns, (b) diet-based inference of personalised gut profiles, and (c) as a steppingstone for studying multi-species metabolic interactions.

## INTRODUCTION

Metabolism in the host is complemented by the microbial community (microbiota) harboured in its gut. The microbiota collectively possesses a larger repertoire of enzymes which helps in digestion and nutrient uptake from sources such as, complex carbohydrates [1]. Microbes also synthesize and make available different key nutrients such as, essential amino acids, vitamins and short chain fatty acids [2,3]. Consequently, imbalances (i.e., dysbiosis) in the gut microbiota impacts an individual’s health and has been linked to many diseases like inflammatory bowel disease, obesity, type II diabetes, etc. [4–8]. Microbiome usually evolves as a complex community [9] and it is imperative to investigate metabolic interconnection and resultant interactions among them. While many microbiome studies derive inferences based on the correlation of abundances (or cooccurrences) of gut microbial species, often so in a disease or a dietary context [10,11], they seldom focus on their ‘metabolic communication’. Deducing such metabolic communications are often cumbersome, time consuming and costly; given that majority of gut micro-organisms are not cultivable under *in-vitro* conditions [12].

Rapid advancement in genome sequencing in recent years have provided new impetus for development of high-quality genome-scale metabolic models which can aid in microbial metabolic network analysis. In addition to the genomic information, these metabolic models can also be adapted to use multi-omics data (viz., proteomics, transcriptomics, metabolomics, etc.) to replicate the metabolic behaviour of an organism under specific environmental conditions, such as nutrient availability, stresses, co-culturing, etc. [13,14]. Earlier works by independent research groups have established that pairwise interactions are the major drivers of bacterial communities, as opposed to their higher◻order interactions [9,15]. Metabolic exchanges between two species could exemplify the nature of interactions that occurs between them [16,17]. This is especially pivotal while considering environmental factors, such as diet which could strongly drive the microbial composition and intrinsic metabolic behaviour inside the gut [18]. Therefore, a joint genome-scale reconstruction of two different organisms, in conjunction with Flux Balance Analysis (FBA) [1,8,16,19–21], could elicit metabolic patterns that would define their innate relationship within a dietary/ nutrient regimen (Dai et al., 2019; Heinken et al., 2019; Perisin & Sund, 2018). This has been famously exemplified by Klitgord and Segrè [23], wherein the authors examined paired combinations of seven metabolically reconstructed microbes (models) to identify nutrient environments that induced symbiotic relations, which would otherwise deter growth in isolated condition. This involved a combinatorial approach in determining media that led to emergent mutualistic dependence through bidirectional exchange of nutrients necessary for growth. It was also surmised that environmental/ nutrient fluctuations could have more profound effect on microbial symbiosis than their genetic (or reactionary) perturbations. Along the same lines, it has been shown that cooperative behaviour occurs when paired-microbes have fewer common growth promoting metabolites [24]. Another study on microbial consortia showed that these pairs/ consortia could produce new metabolites which were otherwise absent in mono-cultures [25]. Some earlier metabolic modelling efforts in this direction have also highlighted the capacity of paired models to produce metabolites which were non-existent in their secluded form, as well as presented examples of the paired models’ increased potential of producing metabolites as compared to the additive sum of the metabolite fluxes in their ‘mono-culture’ simulations [21].

These studies demonstrate the importance of studying interspecies relationships delineating their mutualistic or inhibitory tendencies with each other in a case dependent manner. Our work finds its basis in the above premise, explores the same in context of a human gut habitat, and provides an extensive collection of potential interactions for all gut microbes for which viable metabolic models were available from the VMH (Virtual Metabolic Human) repository (Noronha et al., 2019). The potential interactions are derived from pairwise FBA simulations of gut microbes mimicking their growth in 13 different dietary conditions. Having access to a dietary “interactome”, could provide contextual guidance and justification towards elucidating underlying relations amongst gut microbes, especially so while drawing inference from such relationships determined through microbial abundance-based correlations. Furthermore, one can also posit an approach for delineating key microbial growth deviations within or inter dietary compositions, that would be helpful in understanding individual gut microbiome profiles during a comparative analysis. The pairwise interaction type (as well as growth potential) data for different diet types presented in this work essentially represents a semi-exhaustive collection of gut bacterial ‘dyads’ (the smallest unit of interaction in a social network/ group) and lays the foundation for progressively building onto as well as studying larger gut bacterial networks/ ecosystems.

## RESULTS

Metabolic simulations, based on flux balancing principles, were performed to gauge the growth potential of gut microbes under varying diet conditions. A total of 818 metabolic models resembling human gut associated microbes and 13 diet constraints imitating nutrient availability (to gut microbes) in different dietary habits were used (see MATERIALS AND METHODS). Simulations were performed for single organism models as well as paired organism models to mimic growth of gut microbes in both mono-culture and co-culture conditions under different diet conditions. Further, for each of the diet types, interactions between a pair of microbes was determined from the change in growth rates of the two organisms under co-culture (paired) and mono-culture conditions (see MATERIALS AND METHODS).

### Technical validation against earlier AGORA simulations

The obtained growth rates of the metabolic models representing the gut microbial species, both in mono-culture and co-culture simulations, were benchmarked against the results presented by Magnusdottir et al. [16], who had employed AGORA models (v1.0) in their study. Since their simulation outcomes were reported for only two diet conditions, viz., High-Fiber (AGORA) and Western (AGORA) diet, the evaluation could be performed for these two diets only. For the 768 microbial species (metabolic models) which were common between AGORA (version v1.0) [16] and our present work, we found strong correlation in their single model (mono-culture) growth rates in both High-Fiber (AGORA) as well as Western (AGORA) diets. SRC of 0.921 and 0.954 and PCC of 0.926 and 0.952 were observed for the microbial growth rates in High-Fiber (AGORA) and Western (AGORA) diets respectively. Similarly, comparison of the collective growth rates of the pairwise model (co-culture) also showed good correlations for both the diets (considering 283,881 combinatorial pairs common to both studies). In the co-cultured simulations, SRC of 0.903 and 0.933 and PCC of and 0.87 were noted for High-Fiber (AGORA) and Western (AGORA) diets respectively.

### Assessment of computed interactions in the context of literature evidences

#### Bifidobacterium growth patterns in High Protein and High Fat diets

Using single model simulation results in different VMH Diets, the mean growth rate of 39 different available models of *Bifidobacterium* species was correlated to the main dietary constituents, namely lipids (%), carbohydrates (%), protein (%), dietary fibers (mg), cholesterol (mg) and sugar (mg) (as downloaded from nutrition information table provided in www.vmh.life/#nutrition). Dietary fiber was found to have the strongest positive correlative emergent (PCC of 0.53) of growth rate in single (mono-culture) model condition, and conversely, lipid of the diet showed negative correlation (PCC of −0.49) to growth rate of *Bifidobacterium*. PCCs obtained for the other factors, viz., carbohydrates, protein, cholesterol, and sugar (sucrose) were 0.22, 0.15, −0.19 and 0.24 respectively.

#### Complementarity between Bacteroides thetaiotaomicron and Methanobrevibacter smithii

Two gut inhabiting organisms, *Methanobrevibacter smithii* and *Bacteroides thetaiotaomicron,* are known to exhibit mutualistic (syntrophic) behaviour when grown in a polysaccharide (dietary fiber) based diets [28,29]. We investigated if their syntrophic behavior (in fiber rich diets), could also be replicated in our *in-silico* results. *M. Smithii* (model name Methanobrevibacter_smithii_ATCC_35061) was found to have higher growth rate when co-cultured (paired) with *B. thetaiotaomicron* (model name Bacteroides_thetaiotaomicron_VPI_5482) in fiber rich diets. Its growth rate was seen to increase by 4.51 folds in High-Fiber (AGORA) diet and by 1.44 folds in High-Fiber (VMH) diet. For diets with poor fiber content (like Unhealthy diet and High-Fat with Low-Carb diet), a reverse relationship of amensalism was observed wherein a drop in the growth rate of *M. smithii* by 0.99 folds was found on co-culturing with *B. thetaiotaomicron.*

#### Complementarity between Bifidobacterium adolescentis and Eubacterium hallii

In yet another instance, our simulation results could mimic the commensalistic behaviour between *Eubacterium hallii* (model name Eubacterium_hallii_DSM_3353), a prominent butyate-producing bacterium [30] and *Bifidobacterium adolescentis* (model name Eubacterium_hallii_DSM_3353), in diets which are rich in starch. Notably, it has been reported that *E. hallii* by itself is not able to sustain in a starch rich diet and require assistance from *B. adolescentis* for its survival [4]. In data presented in Table 1 this pair exhibited commensalism in seven out of 13 diets and all of these diets feature higher starch content. Three of the remaining diets (viz., Unhealthy, High-Fiber and Vegan) also had higher starch content, but did not lead to any appreciable increase in the growth of *E. Hallii* (i.e ≥10% of growth rate) and their overall interaction was thus interpreted as neutralism for those diets. Diets with poor starch content yielded negative interactions for this pair.

**Table 1:** Pairwise relationship between *Eubacterium hallii* and *Bifidobacterium adolescentis* under different dietary simulations.

| Interaction | % change in growth<br>rate<br>(*E. hallii*) | % change in growth<br>rate<br>(*B. adolescentis*) | Diet | Starch Uptake<br>(mmol/gDW/hr) |
| --- | --- | --- | --- | --- |
| Amensalism | -61.93 | 0.00 | High-Fat Low-Carb | 0.005 |
| Neutralism | -1.59 | 0.00 | Unhealthy | 3.176 |
| Neutralism | 0.00 | 0.00 | High-Fiber | 3.572 |
| Neutralism | 8.27 | 0.00 | Vegan | 2.444 |
| Commensalism | 33.31 | 0.00 | Vegetarian | 3.273 |
| Commensalism | 47.32 | 0.00 | EU Average | 2.616 |
| Commensalism | 54.08 | 0.00 | High-Protein | 2.145 |
| Commensalism | 66.65 | 0.00 | Type-2 Diabetes | 2.010 |
| Commensalism | 83.11 | 0.00 | Gluten Free | 5.295 |
| Commensalism | 96.92 | 0.00 | Mediterranean | 3.248 |
| Commensalism | 168.93 | 0.00 | DACH | 2.969 |
| Parasitism | 314.82 | -45.55 | Western (Agora) | 0.257 |
| Parasitism | 385.40 | -58.07 | High-Fiber (Agora) | 0.068 |

## DISCUSSION

Genome scale metabolic reconstruction is one of the prime examples of genomics aiding metabolomic research. Continuous growth in this field has propelled the gaining of metabolic insights into complex problems like estimating the growth capacity of a microbe in a nutritional environment [7,23] or cross feeding in a microbial community [17,21]. Hence, a collection of such genome scale metabolic reconstructed models (like VMH repository – www.vmh.life) along with several pre-determined dietary compositions provides an opportunity to compile and build a vast resource of individual and/or symbiotic growth capacity of gut microbes, tailored to these available diets. This, otherwise, via conventional experimental procedures would be cumbersome, time consuming and costly if not infeasible. Here in our study we have computed growths and interactions for 4,182,618 combinations of available microbial pairs, and attempted validation of simulated growth rates and derived symbiotic relationships to existing literature.

The ideal validations for the single model (mono-culture) and pairwise model (co-culture) simulation results would be to compare the *in-silico* results with the experimental growth rates under different diet types. However, given a multitude of factors, including difficulties to replicate the diets in culture media, and challenges in growing most gut microbes in the laboratory, the availability of experimental data to benchmark *in-silico* findings are limited. Consequently, the publication presenting the original AGORA models (v1.0) [16] evaluated simulation results using growth rates of only a single pair of gut microbes under a specialized nutrient environment. This being a seminal publication on the topic, the results presented therein were considered as a benchmark while performing the technical validations for our current study. In brief, the mono-culture and co-culture growth rates of the 773 gut microbial models (from AGORA v1.0), simulated under the two AGORA diets, viz., High-Fiber (AGORA) and Western (AGORA) were used for this comparison. Subsequently, we have also evaluated some of our predicted growth rates and derived symbiotic relationships against experimentally observed diet-linked microbial growths and interaction patterns available from literature.

It may be noted that the current version of AGORA models (v1.03), that has been used for simulations performed in the current study, have been updated and refined since the original publication [16]. The changes include rectification of false positive predictions of nutrient uptakes within the model, implementation of improved gap-filing protocols on a new refined growth media [31], and introduction of new pathway reactions from several studies like aromatic amino acid degradation [32], putrefaction pathways in the gut [33], bile-acid biosynthesis [7]. Given these differences in the models used as well as certain differences in the methodology when compared to Magnusdottir et al. [16], some deviations pertaining to the computed growth values, and the interactions derived, could be anticipated. The methodological differences included usage of some revised reaction constraints (see Diet Construction sub-section of MATERIALS AND METHODS), usage of COBRApy library (python) in place of COBRA toolbox (MATLAB), usage of glpk solver (publicly available) instead of the proprietary CPLEX solver (IBM, Inc.), using an adapted version of Mminte (a python package) for paired model reconstruction [34] (see Code Usage in Appendix 1), and employment of auxiliary flux coupling constraints, implemented within python (see MATERIALS AND METHODS section and Code Usage in Appendix 1). Despite the technical and methodological differences, the two studies displayed similar results in terms of growth rates for the individual and paired organisms (See RESULTS section).

Additional validations were subsequently performed to check if the interaction patterns (and the simulated growth rates) among a pair of microbes, as reported in this work, could replicate the biologically observed phenomenon under different diet conditions. The three case studies (as shown in RESULTS section) highlight the potential use that can be extended in this regard.

Numerous studies have focussed attention to *Bifidobacterium*, an eminent gut inhabiting species, which is particularly known for its probiotic interplay within host and with gut microbial species [35,36]. Studies suggest that *Bifidobacterium* species grows poorly in diet compositions made with high protein [37], and with high-fat and low-carbohydrate [38]. Our simulation data gives similar indications for this species as shown by moderately negative correlation to lipid content (See RESULTS section). It may be mentioned in this context that Hwang and his co-workers [37] also evaluated the growth patterns of *Sutterella*, another gut bacterium, in addition to *Bifidobacterium* and reported contrasting growth trends. Unfortunately, the two models of *Sutterella* which have so far been reconstructed, were a subset of 27 gut bacterial models (out of 803 used in this study) exhibiting no appreciable change in growth rates across diet types and often very poor growth in mono-cultures (Supplementary Table 1 in Appendix 1). Therefore, growth patterns of *Sutterella* in response to different dietary constituents could not be assessed in course of technical validation for this work. While the diet-invariant very low growth rates, possibly due to the inability of these organisms to survive in isolation in the human gut, may be construed as a limitation of this work, it may be noted that the growth rates of these organisms (including *Sutterella*) showed significant variations in the co-culture simulations across different diet types.

Literature evidences also substantiates the simulation results i.e. growth rate derived interaction paradigms, obtained in our study. For instance, *Bacteroides thetaiotaomicron*, one of the most common gut species, and *Methanobrevibacter smithii*, a pre-dominant gut microbe of *Archaea* domain, have been notably shown to have syntrophic relationship, wherein *B. thetaiotaomicron* assists *M. smithii* to grow in polysaccharide (dietary fiber) based diets [28,29]. Aligned with the experimental evidences, we observed commensalism in our paired-model simulations between these two species in diets with high fiber content. On a similar note, our derived interactions between gut microbes could also be validated for another prominent experimental observation [4], which included *Bifidobacterium adolescentis* and *Eubacterium hallii.* From the data presented in Table 1, the above interaction phenomenon could be observed in diets which had higher starch contents, wherein the co-cultured pair tend to display commensalism in favour of *E. hallii*.

Given the above, we believe that the provided resource would be useful in drawing inferences from putative interactions between different gut organisms or from their overall growth patterns across diverse set of pre-determined nutritional compositions. Our simulation data could aid in providing clues (from metabolic perspective) to microbial interrelationships derived solely from abundance-based correlations. And with a wider choice of dietary compositions available to the users, there is an added propensity to mimic the diet of the samples from which those correlations were derived, which makes the inferences/ justifications more meaningful.

## CONCLUSIONS

The datasets generated in our study allows analysis/ data-inferences at intra/ inter diet level, both of which enables investigation of diet induced growth patterns of an organism, a taxonomic group or at the gross level for the entire microbiome samples. This could be useful for investigation/ validation of any symbiotic relationships and growth deviations observed for an organism of interest across single or several diets from experimental or *in-silico* studies. Users can also utilize the pairwise growth values and deploy different growth cut-off parameters for customizing definitions of symbiotic relationships and mining for such interactions in a dataset of interest. In addition, users can make use of the organism’s growth rates/ interaction information for pruning microbial association networks derived from abundance-based studies, as shown in an earlier study [17]. This data makes it possible to filter or validate the edges of interaction networks of gut microbes from abundance-based correlations and justify those connections from metabolic perspective. Furthermore, the scripts provided in the repository allows for the extension of the framework to microbes residing in any ecological niche and is thus expected to be beneficial for microbiologists, ecological experts and other researchers working in allied areas.

## MATERIALS AND METHODS

### Mono-culture (single model) simulation

A total of 818 models representing the metabolic potential of human gut associated microbes were retrieved from AGORA (assembly of gut organisms through reconstruction and analysis) v1.03 (version dated 25-Feb-2019) hosted at www.vmh.life (Noronha et al., 2019) (see Supplementary Table 2 in Appendix 1). While the current version of AGORA metabolic models has been reported to be curated and refined based on experimental evidences in recent scientific publications, for the purpose of the current study, each of the downloaded metabolic models were further modified in the following manner:

a. The reactions and metabolite identifiers within the models were converted to BiGG identifier notation style so as to make it compatible and convenient for its use in with COBRApy package [19].
b. The lower bounds of the exchange reactions were modified to mimic the appropriate diet constraints (see Diet Construction sub-section of MATERIALS AND METHODS). If an exchange reaction of the model was absent in a diet’s constraints list, then the lower bound for that reaction was set to 0.

Finally, FBA was performed on each of the modified metabolic models under different diet constraints (see Diet Construction sub-section of MATERIALS AND METHODS) using glpk solver and COBRApy package in python [19]. The objective of the simulations was to predict maximum possible growth of each of the bacteria (represented by their metabolic models), when grown as anaerobic mono-culture under different diet conditions.

### Co-culture (paired model) simulation

In order to replicate metabolic interactions among a pair of gut microbes, pairwise simulations were carried out for 13 different diets (Table 2). Notably, the metabolic models representing 15 gut microbes showed infeasible FBA solution for growth optimization in at least one of the diets under mono-culture condition and were excluded from the pairwise simulation experiments. All combinations of the remaining 803 models were considered which totalled to 322,003 pairs. The Mminte package [34] in python was employed to reconstruct the paired models (representing a pair of gut microbes) using earlier suggested strategies (Magnúsdóttir et al., 2017; Mendes-Soares et al., 2016). In brief, the models were joined into a common lumen compartment which acted as an extracellular interface for the exchange of metabolites. Additionally, to avoid scenarios where an organism (metabolic model) benefits the other without producing any biomass (i.e. the objective function), flux coupling constraints were introduced which stoichiometrically coupled every reaction to the biomass objective function, as per the strategy suggested in earlier literature [7,16]. After introducing dietary constraints to the extracellular compartment of the model (as followed for single model simulations), FBA was run to simultaneously maximise growth of both organisms. Out of all the 322,003 model pairs, 331 model pairs could not be solved for either one or more VMH diets using glpk solver that was used in this study. The output of each solvable pair, i.e. growth of each organism in paired condition, single condition, percentage growth change between the conditions and finally the interaction type was computed and saved for each diet. Thus, output from 321,692 pairs for each VMH diet and 322,003 pairs for each AGORA diet were tabulated and uploaded to the OSF Home repository.

**TABLE 2:**
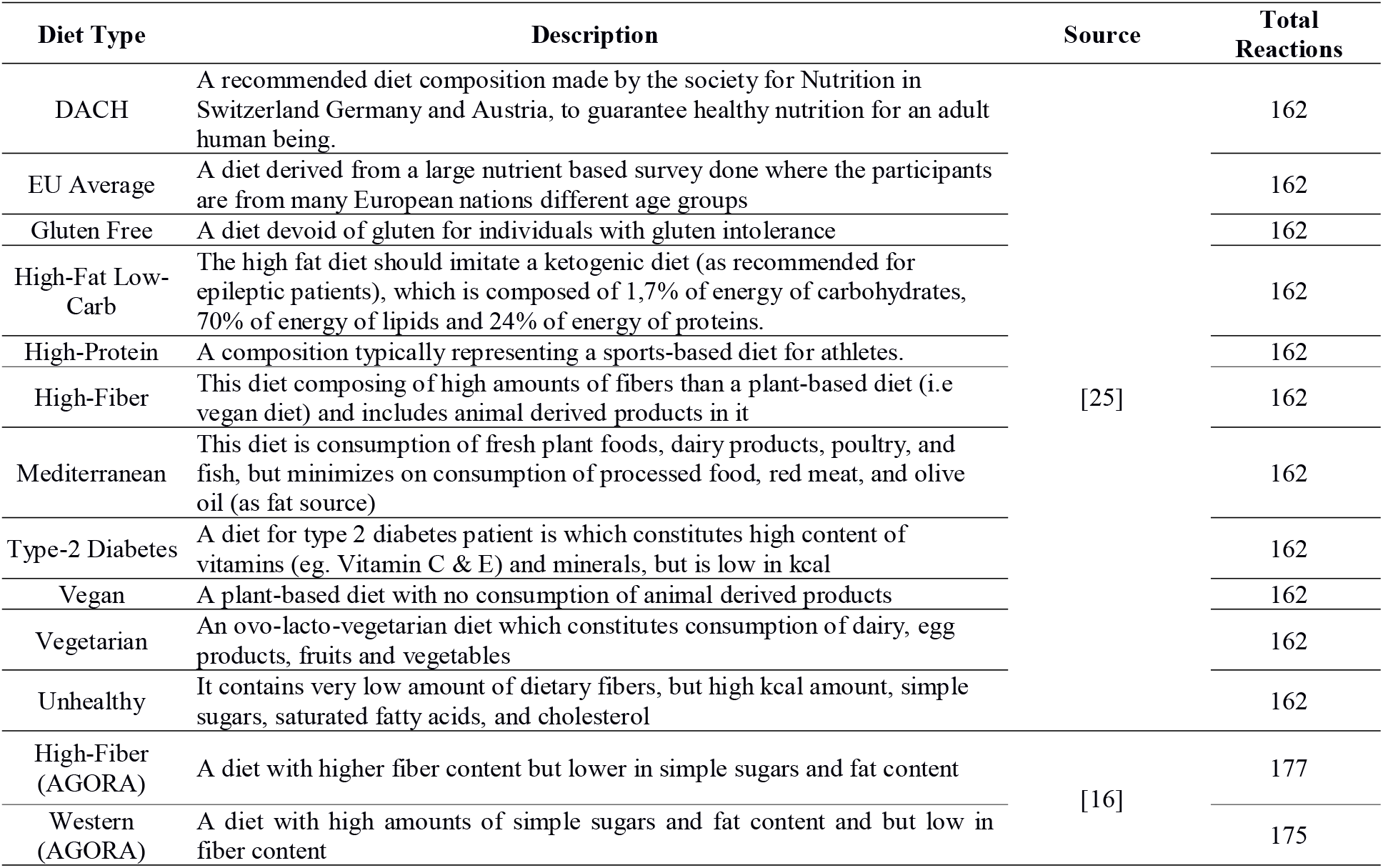
List of the diets used in this study along with the number of their reactionary constraints and the literature where they were first defined.

### Determination of interaction

Interaction types, between each pair of organisms, were evaluated from the simulated growth rates of the organisms under co-culture (paired) and mono-culture conditions (Fig. 1). In line with previous studies [16,21,34], whenever the growth rate of an organism changed by ≥10% during co-culture ([G_org_]^P^), when compared to its growth rate in isolation ([G_org_]^I^), a discernible interaction amounting to a symbiotic relationship was considered (Table 3). Positive influence (+) was denoted for increased growth rate, negative influence (−) for a decrease in growth rate, and no effect (0) if the growth rate did not change by at least 10%. For every given pair of organisms (in a given diet type), one of the six different interactions were assigned based on possible pairwise growth profile outcomes depicted in Table 3.

**TABLE 3:** Pairwise interaction patterns based on the growth profile outcomes of the two organisms constituting a (paired) co-culture simulation experiment.

| Type of Paired Interaction | Description | Abbreviation |
| --- | --- | --- |
| Amensalism | One organism deteriorates in growth while the other organism remains unaffected | (0, -) or (-, 0) |
| Commensalism | One organism increases in growth while the other organism remains unaffected | (0, +) or (+, 0) |
| Competition | Both organisms suffer from drop in their individual growths under paired condition | (-, -) |
| Mutualism | Both organisms have augmentation in their individual growths under paired condition | (+, +) |
| Neutralism | Growths of both organisms remain unchanged under paired condition | (0, 0) |
| Parasitism | Growth of one organism diminishes while the same increases in the other organism | (-, +) or (+, -) |
| **Abbreviation keys -** 0: Unaffected; +: positive change; -: negative change |  |  |

**Figure 1:**
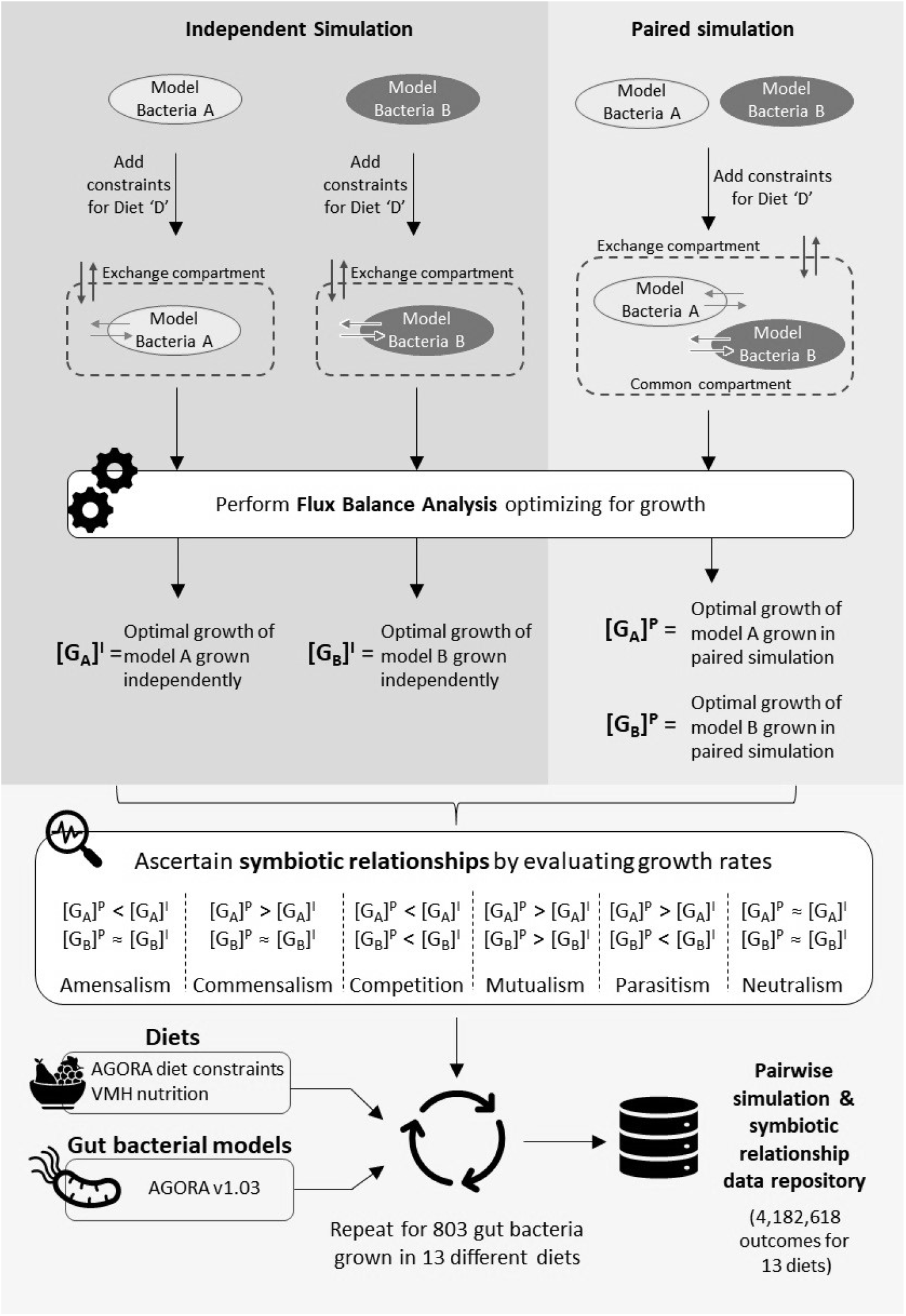
Schematic representation of the process followed for determining pairwise symbiotic interactions between gut microbial species. The ‘>’ and ‘<’ symbols denote that the growth of an organism in paired simulations [G_org_]^P^ (mimicking co-cultures) deviates at least by 10% or more when compared to its growth when simulated independently [G_org_]^I^ (mimicking monoculture).

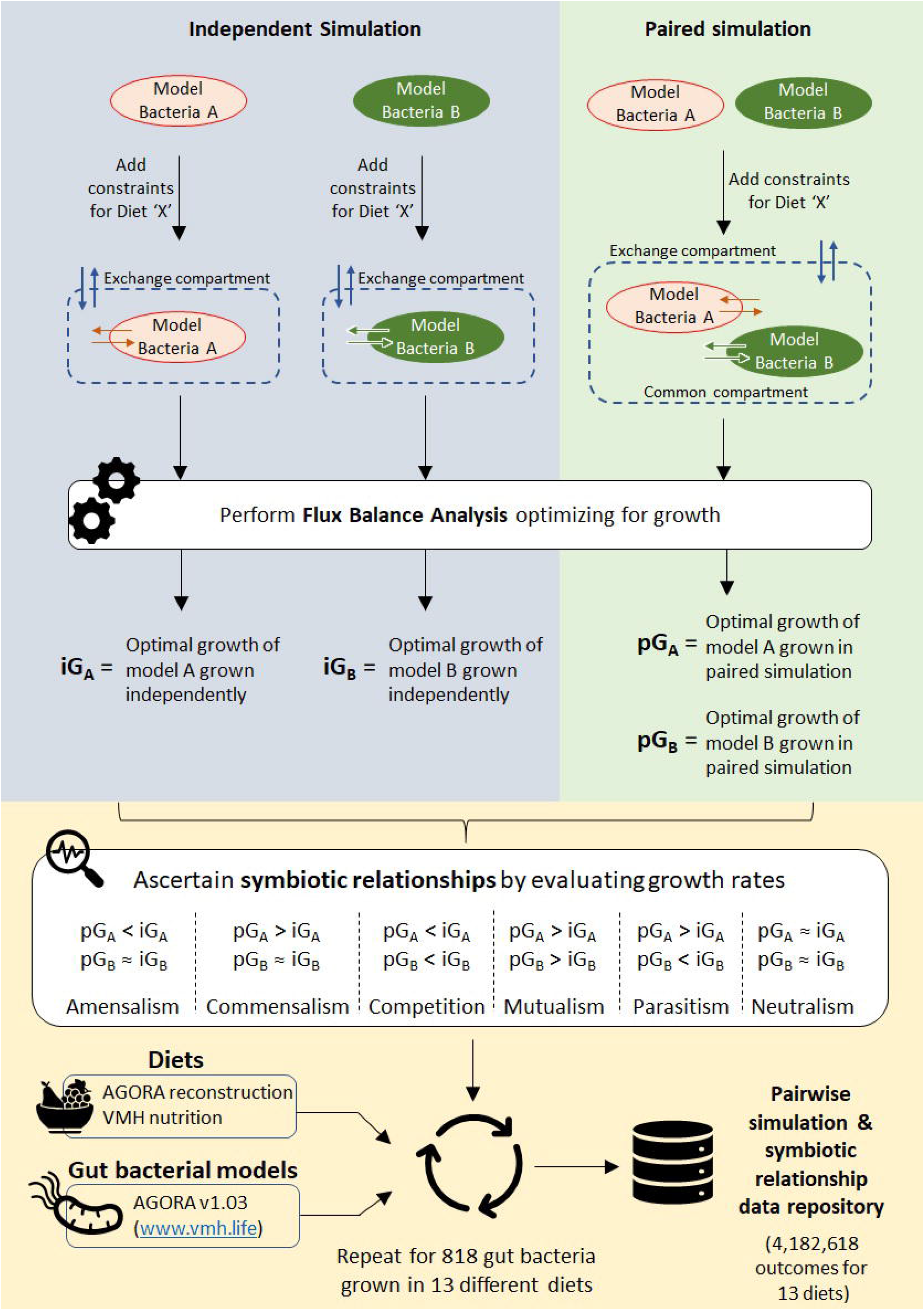

### Diet construction

Human societies around the world have different diet preferences which differ widely in nutrient composition. Gut microbes are known to exhibit alternate metabolic behaviour, and consequently varying growth rates, in response to different diet types [4,18,39]. To mimic this, the metabolic models of the gut microbes were simulated to grow on 13 different diet types (Table 2), as mono- and bi-cultures (paired). Of the total 13 diets used in this study, metabolic exchange constraints representing two diets (High-Fiber and Western) were obtained from Magnusdottir et al. [16]. These two diets were then edited to incorporate modified flux constraints for certain exchange reactions (such as setting lower bounds of exchanges of acetaldehyde, 2-oxoglutarate, L-lactate, L-malate, succinate to 0 mmol/gDW/hr), as mentioned in AGORA v1.01 update (from www.vmh.life). The remaining 11 diets were retrieved from “Nutrition” section of VMH (from www.vmh.life). Since these set of constraints defining the diet types by itself could not support growth for majority of AGORA models, an adaptation protocol was additionally followed (as described in Heinken et al., 2019). This protocol was adapted from “adaptVMHDietToAGORA” functionality of Microbiome Modeling Toolbox [40] and was implemented in python for our study (see Code Usage in Appendix 1).

## Data availability

All data pertaining to this work has been tabulated and archived in OSF Home Data Repository [41]. Details of the data records along with the format for each of the data files are provided in Appendix 1 (see Data Record Information and Supplementary Tables 3, 4).

## AUTHOR CONTRIBUTIONS

R.S., A.D., T.B. and S.S.M. conceived the idea, designed the protocol for data simulation and analysis. R.S. implemented the codes, performed the simulation experiments and created the data repository. R.S., A.D. and T.B. analysed the results. All authors contributed towards drafting the final manuscript.

## DATA AVAILABILITY STATEMENT

The simulation results obtained in this study has been deposited to ‘OSF Home’ repository [41]. The data deposited to ‘OSF Home’ further comprise of the mono-culture and co-culture simulation growth rates of 803 gut microbial species in 13 different diet types and the derived symbiotic relationships between the gut microbial species. Description of the file formats for these data records have been provided in Supplementary Tables 3 and 4 (Appendix 1). In addition, a stand-alone program used for obtaining the co-culture simulation results is also provided. This program accepts, as argument, a pair of genome scale metabolic model files (in json or xml format) and a diet file (in json format) to generates co-culture growth rates of the two microbes as well as infer the type of interaction among them. All these resources are freely accessible for academic use.

## ABBREVIATIONS USED

FBA: Flux Balance Analysis
VMH: Virtual Metabolic Human
AGORA: Assembly of Gut Organisms through Reconstruction and Analysis
SRC: Spearman’s Rank Correlation Test scores
PCC: Pearson Correlation Coefficients

