## Appendix 1 for "A compendium of predicted growths and derived symbiotic relationships between 803 gut microbes in 13 different diets"

**A compendium of predicted growths and derived symbiotic relationships between 803 gut microbes in 13 different diets**

Rohan Singh, Anirban Dutta, Tungadri Bose*, Sharmila S. Mande*

TCS Research, Tata Consultancy Services Ltd., 54-B Hadapsar Industrial Estate, Pune, 411 013, India

*Corresponding authors

**Data Record Information**

***1) Data Summary***

As has been presented in Supplementary Table 1 the simulated growth rates of 803 gut bacterial species under 13 different diet types have been provided ((***Mono-culture Simulation Data)***. The metabolic models as well as the diet constraints (***Diet Constraint Data***) were obtained from AGORA Reconstructions (Magnúsdóttir et al. 2017) and VMH Nutrition (Noronha et al. 2019). Further the growth rates and symbiotic patterns under (paired) co-culture conditions have been provided (***Co-culture Simulation Data***). Based on the change in growth rates under co-cultured conditions, the interaction patterns between a pair of organisms have been bracketed into six categories (see section 2.3 of MATERIALS AND METHODS section).

It may be noted that each entry within the co-culture simulation dataset is a unique combination, i.e. for a pair of organisms A and B, if A-B model information is present, then B-A combination is considered a duplicate and will not be present.

***2) Diet Constraint Data***

All the 13 diet constraints have been provided as individual JSON files. Each diet file includes “keys” and their associated “values”. Here, each key refers to the name of the exchange reaction (in BiGG identifier notation format) corresponding to a dietary component, and the value of each key is the constraint value (lower bound) for that exchange reaction.

***3) Mono-culture Simulation Data***

Data pertaining to mono-culture (single model) simulations has been documented in a comma separated text (csv) file, and uploaded to the aforementioned repository. This file consists of growth rates of each organism used in this study for all 13 diet types. The description of each of the columns in the csv formatted file is provided in Supplementary Table 2.

***4) Co-culture Simulation Dataset***

This data has been presented in comma separated text (csv) files, corresponding to the respective diets. Each entry within the data file corresponds to a pair of gut micro-organisms, containing information about their names (at strain level), the combined growth rate, their respective growth rates in single and paired conditions for the diet, their percentage change(s) in growth and the symbiotic relationship evaluated for that pair (Supplementary Table 3). Name of the csv files is indicative of the respective diet types (e.g. Pairwise_simulation_data-Vegan.csv correspond to Vegan diet).

**Code Usage**

For the purpose to recreate our datasets or to construct and evaluate new paired models for different datasets, python scripts have been provided. All the paired model and result generation were done by the following scripts:

***1) pair_FuncDefinitions.py***

Includes python function definitions derived from Mminte package for model creation, our custom-built functions for diet and flux-coupling constraints integration, and cut-off parameters for evaluating interactions and growth optimality status.

***2) pair_Main.py***

A python3 executable script, to which models and diet files are provided as positional arguments. It takes two model files (in json or xml format) as arguments followed by a diet file (in json format). This script file calls the pair_FunctDefiniation.py when executed, hence these two script files need to be placed within the same folder to generate the output.

Users can run the code in the folder containing the gut models and a diet constraints file. To call the script, run the following command:

python pair_Main.py model_A.xml model_B.xml diet.json

This yields growth rates and interaction output in the form described in Supplementary Table 3 and saves to file “results.csv”. Users can also customise the code to save the models for further analysis.

***3) adaptVMHDietToAGORA.py***

Converts and adapts a VMH diet in a form which makes AGORA models conducive in that diet (refer Methods under ‘Diet Construction’ for further details). This script takes an argument which is a tab separated reaction flux file (as downloaded from https://www.vmh.life/#nutrition) for a VMH diet. To use the script, run the following command:

python adaptVMHDietToAGORA.py fluxes.txt

This will generate a json file “vmhToAGORA_diet.json” which can be used with pair_Main.py (as described above).

**Supplementary Table 1:** Summary of the datasets used, and simulations performed in the presented work.

| **Category** | **VMH Diets** | **AGORA Diets** |
| --- | --- | --- |
| Total Diets | 11 | 2 |
| Total Pairs | 3,538,612 | 644,006 |
| Total Size | 765 MB | 140 MB |
| **Category** | **Number** |  |
| Total Models Used (starting point) | 818 |  |
| Total Models Simulated | 803 |  |
| Interaction Types | 6 |  |
| Nutritional Information Resources Used | 2 |  |

**Supplementary Table 2:** Description of each column in the csv file enlisting the mono-culture (single model) simulation results.

| **Column Name** | **Data Format** | **Column Description** |
| --- | --- | --- |
| Organism | Character | Name of the organism model (sbml model filename) |
| High-Protein, .....,Western (AGORA) | Numeric | Growth of the organism in the diets as indicated by the name of the respective column |

**Supplementary Table 3:** Description of each column in the csv file enlisting the co-culture (pairwise) simulation results.

| **Column Name** | **Data Format** | **Column Description** |
| --- | --- | --- |
| Org1 | Character | Name of the first organism model (sbml model filename) |
| Org2 | Character | Name of the second organism model (sbml model filename) |
| Interaction | Character | Type of interaction deduced (See MATERIALS AND METHODS for details) |
| Together | Numeric | Sum of growth of both organisms under paired form |
| Org1_Paired | Numeric | Growth of first organism in paired condition |
| Org2_Paired | Numeric | Growth of second organism in paired condition |
| Org1_Single | Numeric | Growth of first organism in isolated condition |
| Org2_Single | Numeric | Growth of second organism in isolated condition |
| Change%_Org1 | Numeric | Percentage change between growth of first organism in paired condition when compared to its growth rate in isolated condition |
| Change%_Org2 | Numeric | Percentage change between growth of second organism in paired condition when compared to its growth rate in isolated condition |
| Diet | Character | Name of diet used |

**Supplementary Table 4:** Metabolic models representing gut bacterial strains whose growth rates did not change across different diets. The models for *Sutterella* are marked in bold font.

| **Gut bacterial strains**  **(sbml model filenames)** | **Mean growth rate across diets (mmol/gDW/hr)** | **Standard deviation in growth rate across diets** |
| --- | --- | --- |
| Acidaminococcus_intestini_RyC_MR95.xml | 0.166396826 | 5.01E-15 |
| Adlercreutzia_equolifaciens_DSM_19450.xml | 0.001722732 | 6.82E-19 |
| Bacteroides_oleiciplenus_YIT_12058.xml | 0.305825361 | 6.18E-16 |
| Bartonella_quintana_Toulouse.xml | 0.002229275 | 5.99E-05 |
| Bilophila_wadsworthia_3_1_6.xml | 0.12533524 | 8.05E-16 |
| Burkholderiales_bacterium_1_1_47.xml | 0.000876375 | 3.41E-19 |
| Campylobacter_gracilis_RM3268.xml | 0.001450117 | 2.27E-19 |
| Campylobacter_hominis_ATCC_BAA_381.xml | 0.002821385 | 4.55E-19 |
| Campylobacter_rectus_RM3267.xml | 0.000880091 | 0 |
| Campylobacter_showae_CSUNSWCD.xml | 0.000918164 | 1.14E-19 |
| Cloacibacillus_evryensis_DSM_19522.xml | 0.182836247 | 1.42E-15 |
| Clostridium_sporogenes_ATCC_15579.xml | 0.000225 | 6.41E-20 |
| Corynebacterium_propinquum_DSM_44285.xml | 0.011416223 | 0 |
| Dialister_invisus_DSM_15470.xml | 0.000959435 | 1.14E-19 |
| Faecalibacterium_prausnitzii_M21_2.xml | 0.000225 | 2.84E-20 |
| Faecalibacterium_prausnitzii_SL3_3.xml | 0.000225 | 7.52E-20 |
| Helicobacter_winghamensis_ATCC_BAA_430.xml | 0.012975042 | 1.82E-18 |
| Lactobacillus_coleohominis_101_4_CHN.xml | 0.092362105 | 0.000510785 |
| Lactobacillus_curvatus_CRL_705.xml | 0.092362105 | 0.000510785 |
| Mogibacterium_timidum_ATCC_33093.xml | 0.001091759 | 0 |
| Mycoplasma_hominis_ATCC_23114.xml | 0.001601601 | 0 |
| Neisseria_cinerea_ATCC_14685.xml | 0.405350628 | 1.01E-15 |
| Slackia_exigua_ATCC_700122.xml | 0.039089145 | 1.73E-16 |
| Streptococcus_anginosus_1_2_62CV.xml | 0.086689668 | 3.18E-16 |
| **Sutterella_parvirubra_YIT_11816.xml** | **0.001048941** | **2.27E-19** |
| **Sutterella_wadsworthensis_3_1_45B.xml** | **0.008394685** | **1.82E-18** |
| Tropheryma_whipplei_str_Twist.xml | 0.000881657 | 1.14E-19 |
| Ureaplasma_parvum_serovar_1_str_ATCC_27813.xml | 0.000573899 | 1.14E-19 |
| Ureaplasma_urealyticum_serovar_8_str_ATCC_27618.xml | 0.004325956 | 0 |
